# The untapped potential of conservation journals to promote freshwater biodiversity

**DOI:** 10.1101/2020.07.21.214288

**Authors:** Fengzhi He, Sonja C. Jähnig, Annett Wetzig, Simone D. Langhans

## Abstract

Freshwater ecosystems are amongst the most diverse ecosystems on the planet. They are subject to intense and increasing threats and have a higher proportion of threatened and extinct species than terrestrial or marine realms. Concurrently, freshwater ecosystems are largely underrepresented in both conservation research and actions arguably as a consequence of less popularity and promotion. To test this assumption, we used cover images as a proxy of exposure and promotion opportunities provided by conservation journals. We collected information on cover images of 18 conservation journals from 1997 to 2016 and data on citations and Altmetric scores of papers published in them. We found that freshwater ecosystems (10.4%) were featured less often than marine (15.2%) or terrestrial (74.4%) ecosystems on covers of these journals. All 15 most featured species are from terrestrial or marine ecosystems, with 14 of them being large vertebrates such as elephants, big cats, rhinos, polar bears, and marine turtles. None of the 95 species featured more than once on the covers of conservation journals spend their whole life history in fresh waters, i.e. they are at least partly associated with terrestrial or marine ecosystems. Our results indicated that cover-featured studies received more attention from academia and the general public, i.e. showed higher citations and Altmetric scores, than non-featured ones within the same issue. By featuring freshwater species and habitats on covers, therewith providing more exposure opportunities, conservation journals hold the potential to promote biodiversity conservation in fresh waters. Scientists can help that endeavour by submitting freshwater-related photos together with their manuscripts for review, therewith providing more options for editors to portray freshwater species and habitats and to ultimately raise awareness and appreciation of freshwater life.

## INTRODUCTION

Freshwater habitats including rivers, lakes, and wetlands cover less than 3% of Earth’s surface but support approximately 9.5% of all described animals and one-third of vertebrates (Balian et al., 2008). Meanwhile, freshwater ecosystems are subject to tremendous and increasing pressures due to a growing demand for water, energy, and food, leading to overexploitation of freshwater and organisms (Dudgeon et al., 2006; Vörösmarty et al., 2010; Reid et al., 2019), and to the loss of important habitats such as wetlands and free-flowing rivers (Reis et al., 2017; Grill et al., 2019). Consequently, 27% of all assessed freshwater species are considered as threatened with extinction on the International Union for Conservation of Nature (IUCN) Red List of Threatened Species (Tickner et al., 2020), while global freshwater vertebrate populations have declined by 83% from 1970 to 2014 (WWF, 2018).

Although the proportions of threatened and extinct species and the decline rate of vertebrate populations are much higher in fresh waters than those in terrestrial or marine ecosystems (Costello, 2015; McRae et al., 2017), freshwater ecosystems are largely underrepresented in biodiversity research and conservation actions (Kalinkat et al., 2017; Jucker et al., 2018; Mazor et al., 2018; Tydecks et al., 2018). Even more worryingly, gaps in conservation actions could be worse than those in research (Clark & May, 2002). Indeed, globally 89% of seasonal freshwater wetlands are not covered by protected areas (Reis et al., 2017), and most of the world’s largest rivers have less than 10% of their basins targeted by integrated protection, which falls short of the target (i.e. 17%) of the Convention on Biological Diversity (Abell et al., 2017). Even within protected areas, stressors to freshwater biodiversity often exist. For example, over 1200 large dams and 500 proposed hydropower dams are located within protected areas, which affects the effectiveness of the protection of freshwater ecosystems (Thieme et al., 2020).

Research and conservation actions to safeguard freshwater biodiversity are likely inadequate as a consequence of low popularity among the general public (Monroe et al., 2009; Cooke et al., 2013). Unlike terrestrial and marine ecosystems represented by popular species such as the big cats, elephants, rhinos, polar bears, and cetaceans, freshwater life remains inconspicuous to the public eye and consequently out of sight and out of mind (Monroe et al., 2009; Darwall et al., 2018; He & Jähnig, 2019). Indeed, public perception and knowledge on biodiversity, including its status and importance, are influenced by available information (Papworth et al., 2015; Kochalski et al., 2019), which is currently biased towards certain species and ecosystems (Clark & May, 2002; Jucker et al., 2018; Mazor et al., 2018; Tydecks et al., 2018).

One of the common practices to increase public awareness on biodiversity is featuring species or habitats that are in need of conservation (Clucas et al., 2008). Within the scientific community this is commonly done by conservation journal using species or habitat images as journal covers to promote content, relating the cover image to one of the articles published in the same issue (e.g. Conservation Biology, Diversity and Distributions, and Ecography). These featured species and habitats and related articles are often promoted by journals on social media, which has become an important platform for communicating science and promoting biodiversity conservation (Parsons et al., 2014; Bombaci et al., 2016; Lamb et al., 2018). More and more scientists, conservation journals (e.g. Conservation Biology, Conservation Letters, Animal Conservation, and Ecography) and conservation organizations (IUCN, WWF, Conservation International, and The Nature Conservancy) are active on social media platforms such as Twitter and Facebook, and frequently interact with the general public through these channels (Parsons et al., 2014).

Here we explored the idea that there is an untapped potential of conservation journals to promote freshwater biodiversity by providing more exposure opportunities. Since previous studies have suggested that freshwater ecosystems received less attention from biodiversity research and conservation efforts than terrestrial or marine ecosystems (Jucker et al., 2018; Mazor et al., 2018; Tydecks et al., 2018), we, first, hypothesized that freshwater species and habitats are featured less often on covers than terrestrial or marine ones. Second, we hypothesized that cover-featured articles can reach a broader audience and, therefore, receive more attention in the scientific community as well as generally in society than the non-featured articles within the same issue. If these two hypotheses hold true, freshwater biodiversity could benefit from more exposure opportunities for freshwater studies and related cover images, with likely further-reaching benefits for their protection.

## METHODS

To test our hypotheses, information on cover images of conservation journals from 1997 to 2016 was collected. There are 56 academic journals listed under the category of “biodiversity conservation” in Web of Science database. Among these journals, 18 journals were selected as they regularly changed their covers between 1997 and 2016 and had information on their covers available online or in the printed copies (Table S1). For each cover image, information on the species or habitats featured on cover images was collected. In total, 1043 images with a clear focus on species or habitats and associated ecosystems were included in our analysis. In addition, information on locations (i.e. country or region where photos were taken) was gathered, if it was indicated. When a species was assessed by the IUCN Red List (IUCN, 2018), its associated ecosystems were assigned following the IUCN Red List. For species which are not on the IUCN Red List, a single ecosystem (i.e. freshwater, marine, or terrestrial) or a combination of ecosystems (e.g. marine and terrestrial) was assigned, according to their life history. Similarly, covers that featured habitats only (without species) were either assigned to a single ecosystem or a combination of ecosystems. In case of multiple ecosystems featured on the same cover, the cover count was split proportionally (e.g. 0.5 points for the terrestrial and the freshwater ecosystem count, if both are shown on the cover).

Citation was used as a proxy to measure attention received by published articles from academia. In addition, the Altmetric score was chosen as an indicator of attention from both scientists and the general public. The Altmetric score is a web-driven metric capturing coverage and mentionings on web-based media including news, blogs, social media, and policy documents (Costas et al., 2015). It is considered as a complementary metric to citations, as it can capture broader attention from both scientists and the general public (Piwowar et al., 2013; Bornmann et al., 2014).

For nine journals including Animal Conservation, Conservation Biology, Conservation Letters, Diversity and Distributions, Ecography, Global Change Biology, Journal of Applied Ecology, Oryx, and Systematic and Biodiversity, cover images are usually related to articles within the same issue. Citations and Altmetric scores of articles (excluding editorials and book reviews) published in these nine journals between 2014 and 2016 was collated. The citations of articles were derived from Web of Science on October 27^th^, 2017. The Altmetric scores were collected from journal websites. Considering the fact that Altmetric scores could change over time, Altmetric scores of articles published in the same issue were collected on the same day. Then the percentiles of cover-featured articles within the same issue were calculated, for both citations and Altmetric scores. Wilcoxon signed-rank test was used to test if the percentiles of cover-featured articles are higher than the median (i.e. Q_50_).

## RESULTS

In total, 74.4% of all cover images were related to terrestrial ecosystems, outnumbering the sum of cover images featuring marine (15.2%) or freshwater (10.4%) ecosystems. From 1997 to 2016, terrestrial species and habitats constantly dominated covers of conservation journals (Figure 1), contributing at least 70% to all cover images in each year except for 2010 (64.2%). Since 2007, freshwater ecosystems have been portrayed the fewest in each year. Species and habitats in the USA contributed the highest number (210) of cover images (Figure 2), followed by Canada (26), Brazil (18), Australia (15), and South Africa (15).

**Figure 1.**
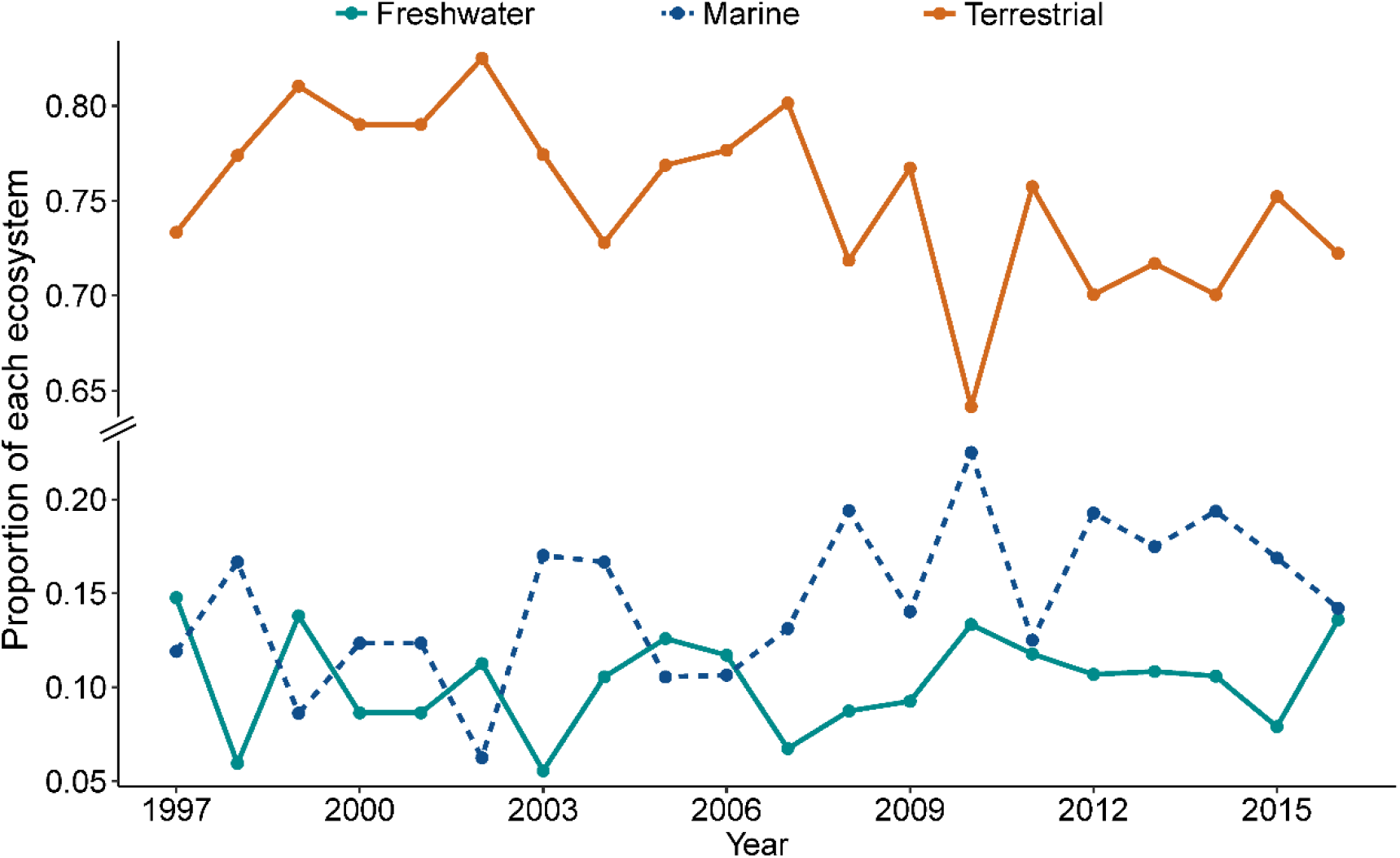
Proportion of species and habitats from each ecosystem on the covers of 18 conservation journals between 1997 and 2016.

**Figure 2.**
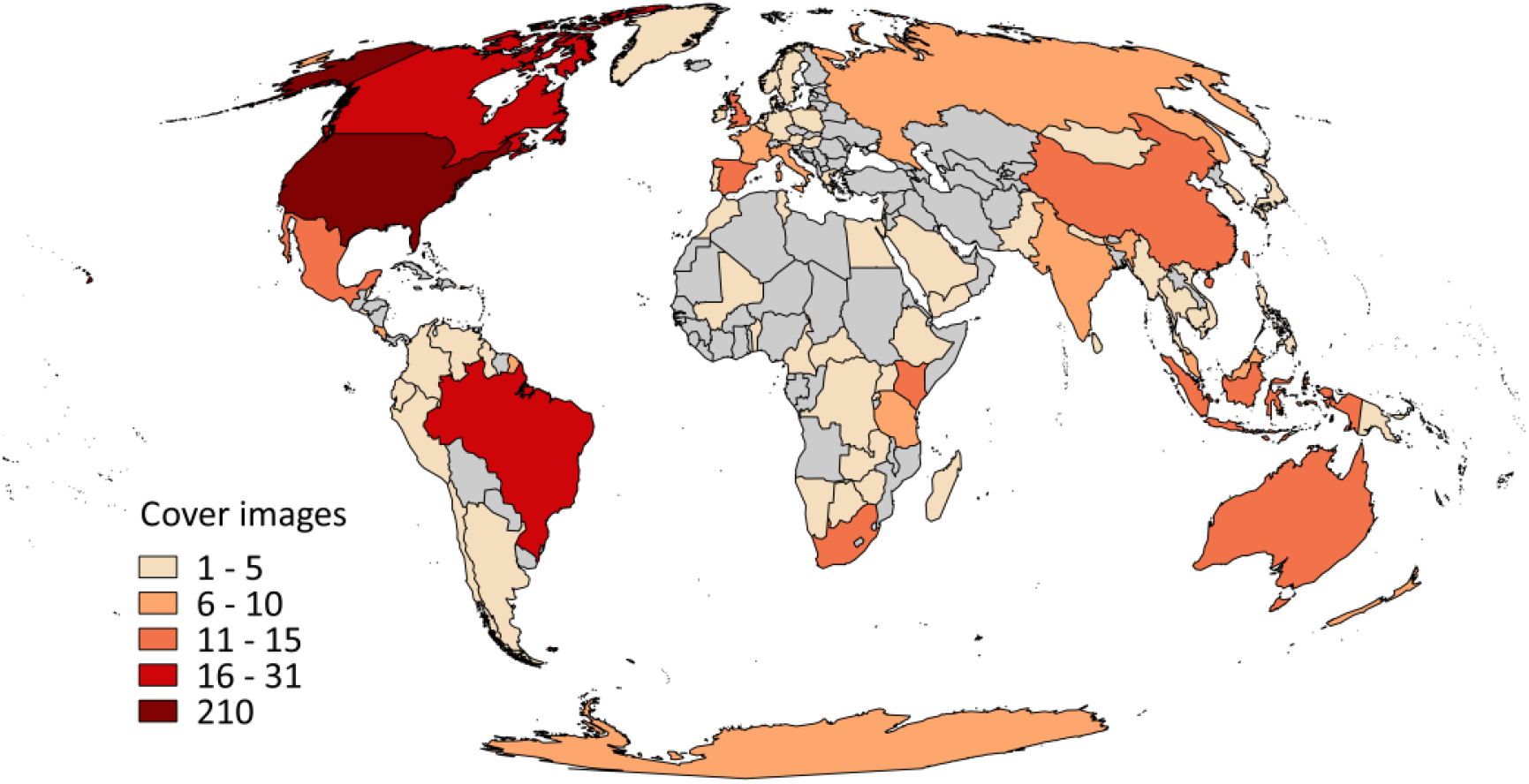
Number of cover images of 18 conservation journals taken in different countries/regions between 1997 and 2016.

In terms of individual species (Figure 3), the African elephant (*Loxodonta Africana*; 18 times) was featured most often on journals’ covers, followed by the tiger (*Panthera tigris*; 8), the black rhinoceros (*Diceros bicornis*; 8), the polar bear (*Ursus maritimus*; 7), the puma (*Puma concolor*; 7), the gray wolf (*Canis lupus*; 6), and the American black bear (*Ursus americanus*; 6). All 15 most featured species (i.e. featured on journal covers at least 4 times) were from terrestrial or marine ecosystems. Fourteen of them are large vertebrate species with the monarch butterfly (*Danaus plexippus*) being the only invertebrate species.

**Figure 3.**
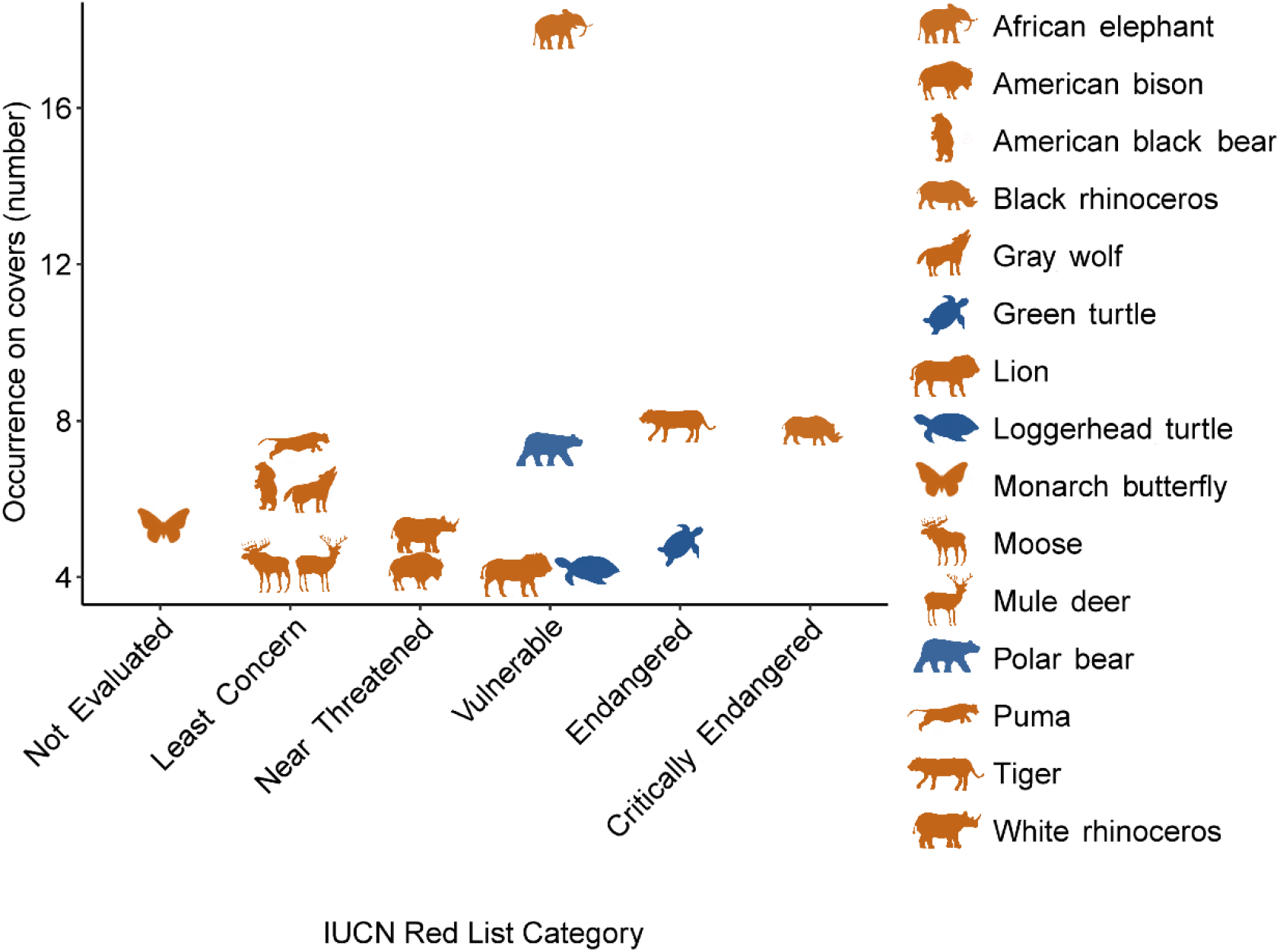
The 15 most featured species and their IUCN Red List categories on covers of 18 conservation journals between 1997 and 2016 (IUCN, 2018; brown-colored animals are from terrestrial ecosystem while blue-colored animals are associated with both marine and terrestrial ecosystems).

Among the 34 species that were featured at least 3 times on journal covers, only 3 species were associated with fresh waters (i.e. *Alligator mississippiensis*, *Ambystoma maculatum*, and *Oncorhynchus nerka*), while 6 species were associated with marine and 32 species with terrestrial ecosystems. None of the 95 species featured more than once was solely associated with fresh waters. Meanwhile, 6 of them were only associated with marine ecosystems and 62 species were only associated with terrestrial ecosystems.

The median percentiles of citations and Altmetric scores of cover-featured articles were 0.63 and 0.76, respectively (Figure 4). The Wilcoxon signed-rank test showed that featured articles had significantly higher citations (p < 0.001, effect size = 0.34) and Altmetric scores (p < 0.001, effect size = 0.60) than non-featured ones within the same issue.

**Figure 4.**
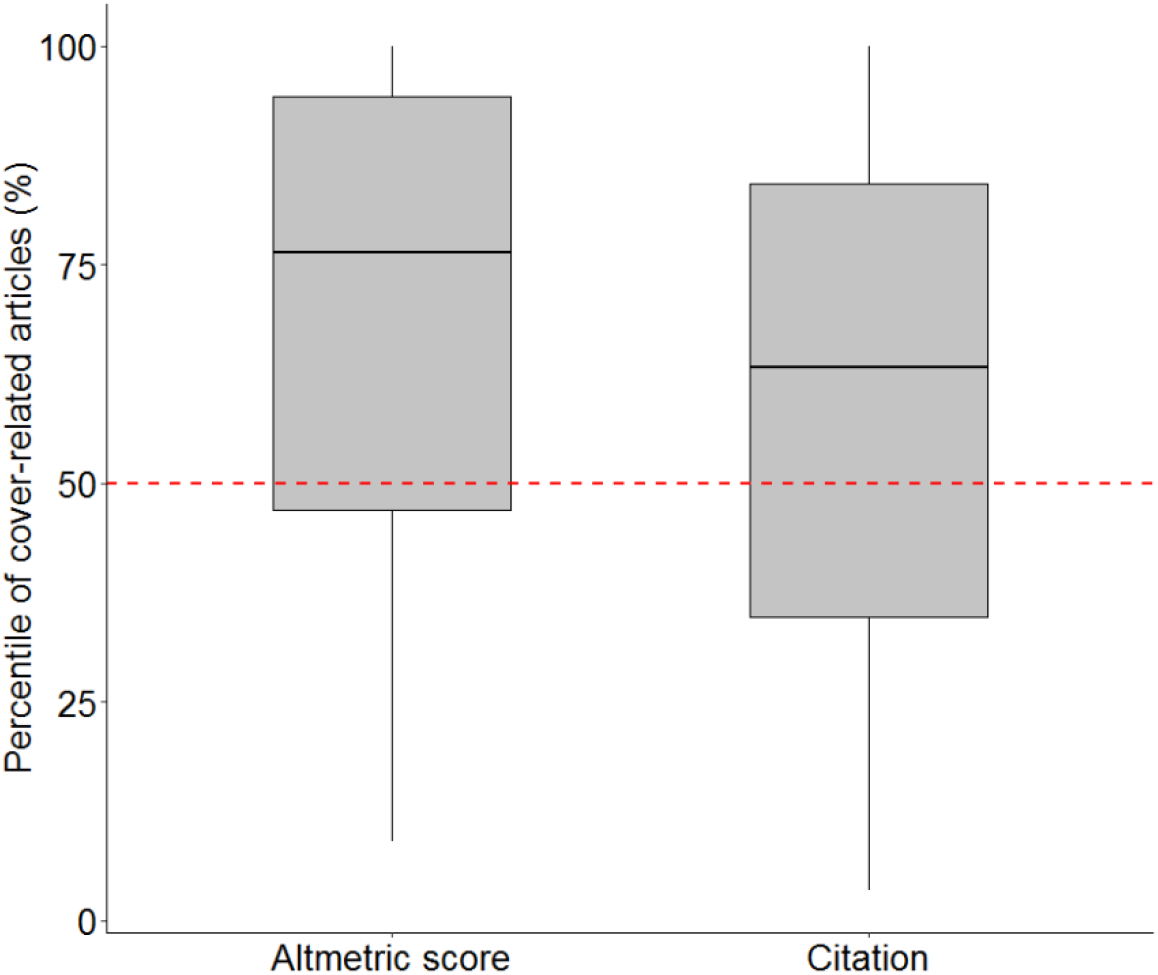
Percentiles of cover-related articles in terms of Altmetric scores and citations within respective issues (N = 168). Red dash line shows the median percentile (i.e. Q50) of all articles within the same issue.

## DISCUSSION

Our results showed that the distribution of cover images and the related, featured studies were skewed in terms of geographical region and ecosystem. Regions harboring a high amount of biodiversity such as Central Africa, Central America, and South and Southeast Asia are currently vastly underrepresented on journal covers. This supports previous findings that most research and conservation efforts have not focused on the regions where they are most needed (Wilson et al., 2016; Tydecks et al., 2018). In addition, we found freshwater species and habitats being largely underrepresented on covers of conservation journals. Our results are consistent with the findings of Clucas et al. (2008) who found big cats, bears, primates, and large birds are often featured on covers of popular conservation and nature magazines in the USA, while freshwater species such as fish and amphibians were rarely featured. Hence, the covers of conservation journals reflect the current research landscape of biodiversity conservation; so far, most research and conservation efforts have focused on terrestrial and marine ecosystems, particularly on large vertebrates (Clucas et al., 2008; Mazor et al., 2018; Tydecks et al., 2018), while only 18% of all biodiversity studies published from 1945 to 2014 are associated with freshwater ecosystems (Tydecks et al., 2018). This is despite that an urgent need for the conservation of freshwater ecosystems has been addressed over 15 years ago (Abell 2002) and a large body of research shows that threats to freshwater habitats and species are intense and increasing over the last few decades (Dudgeon et al., 2006; Vörösmarty et al., 2010; Reid et al., 2019; He et al., 2018; Grill et al., 2019). Terrestrial and marine megafauna species, which are frequently featured on covers of conservation journals (Figure 3) and popular conservation and nature magazines (Clucas et al., 2008), are the ones that receive most of research and conservation efforts (Donaldson et al., 2016; Ford et al., 2017). These species are also the ones on the list of the 10 most charismatic animals perceived by the general public (Courchamp et al., 2018), while freshwater megafauna are often overlooked (Cooke et al., 2013; Carrizo et al., 2017; He & Jähnig, 2019). Tellingly, no freshwater species has made it onto this list.

Three factors may have contributed to the underrepresentation of freshwater species and habitats on the covers of conservation journals: First, fresh waters are often regarded as a resource rather than as important ecosystems. This is despite the fact that they harbor a high level of biodiversity (e.g. 126, 000 animal species), while providing vital ecosystem services (Postel & Carpenter 1997; Balian et al., 2008). Second, compared to terrestrial species, photographers less often portray freshwater species in their natural habitats, but instead display them as “fish out of the water” (i.e. fish species as food or trophy of angling games; Monroe et al., 2009). In addition, large rivers are often turbid, which makes it challenging to photograph freshwater life and underwater habitats compared to marine species and environments. Third, there are generally fewer freshwater studies published in biodiversity and conservation journals than studies focusing on terrestrial or marine ecosystems (Mazor et al., 2018; Tydecks et al., 2018), which therefore limits the choices for editors to display freshwater-related cover images.

Popularity of species or ecosystems can also be generated through global initiatives, such as the Census of Marine Life, which has major positive effects on both public perception and conservation actions (Williams et al., 2010; Vermeulen 2013). So far such prominent, large-scale projects are lacking for fresh waters. In addition, charismatic flagship species are much less promoted in fresh waters in comparison with marine ecosystems that are well represented by popular taxa such as whales, dolphins, sharks, and polar bear (Cooke et al., 2013; Kalinkat et al., 2017; Carrizo et al., 2017; He et al., 2018).

Within the same journal issue, we found that cover-featured articles have higher citations and Altmetric scores than non-featured ones. This indicates that articles featured on covers received more attention from scientists and the general public. On the one hand, such a correlation does not necessarily imply a causation, i.e. the high citations and Altmetric scores of cover-featured articles may not solely be a result of being promoted on journal covers. It is possible that these cover-featured articles received more attention just because they are more interesting and attractive to scientists and the general public than non-featured ones. As experienced scientists, editors often have a good instinct in selecting potentially popular studies that resonate well in academia and the society. In this case, our results only verified good decisions of editors but not the power of cover images. To this argument adds that nowadays the majority of journal readers access research articles through online portals rather than reading the printed copy, and therefore do not come across the journals’ covers. On the other hand, being featured on journal covers can offer more opportunities of exposure to potential readers. For example, cover images are often displayed in prominent positions on websites of conservation journals (e.g. Diversity and Distribution, Animal Conservation, and Journal of Applied Ecology) and are specially mentioned by the journals’ accounts on social media platforms together with the featured study. Cover-featured articles are more likely to be noticed, spread through the internet, and picked up by media outlets, and are, in turn, exposed to a more diverse, non-scientific audience than non-featured articles (Lamb et al., 2018). Hence, the selection of cover images and related featured research by conservation journals entails the potential to facilitate and balance the development of conservation actions.

Editors may be limited in their options when it comes to the selection of a cover image. For example, many papers only show figures of data and model results which do not provide attractive journal covers. In addition, fewer freshwater studies are accepted in biodiversity and conservation journals than marine or terrestrial ecosystems (Mazor et al., 2018; Tydecks et al., 2018), which makes it challenging for editors to balance the journal covers among ecosystems. By submitting appealing images of freshwater species and habitats to journals as potential cover images along with their freshwater-related articles, scientists can play an active role to support editors in promoting freshwater research. Such images can also be used by journals to promote articles on social media platforms. By doing so, scientists and editors can form an alliance to create momentum in society for fresh waters to be experienced as essential ecosystems harboring charismatic species and providing important ecosystem services. Moreover, scientists can directly enhance their communications with decision makers, media, and the general public to inform them about the need of biodiversity conservation in fresh waters. Such direct interaction has been suggested to influence conservation actions (Parsons et al., 2014; Papworth et al., 2015) and lead to better uptake of science in policy (King et al., 2017).

Studies that make it into one of the conservation journals are all significant and novel and have, therefore, a certain potential to be featured on covers. Thus, conservation journals could work towards balancing their choice, inviting more freshwater scientists as editors and providing more exposure opportunities for freshwater-related studies, whenever such an opportunity arises. To support conservation journals in this endeavor, we encourage scientists to include their favorite freshwater photos in future manuscript submissions. We also call for scientists, conservation organizations, and photographers to work together to portray more freshwater species and habitats, raising public awareness and appreciation of freshwater life. In addition, more studies are needed to explore the roles of conservation journals and their social media accounts in promoting biodiversity conservation. For example, it would be interesting to examine the proportion of scientists versus non-scientists in their followers and what makes a post to be retweeted, liked and commented on by these followers. Hence, this study allows formulating specific hypotheses to be tested in future studies, which is a necessary step in solving the major task of safeguarding freshwater ecosystems and its biodiversity that lies ahead of us.

## ACKNOWLEDGEMENTS

This work has been carried out within the SMART Joint Doctorate (Science for the MAnagement of Rivers and their Tidal systems), funded with the support of the Erasmus Mundus programme of the European Union and is a contribution to the Leibniz Competition project “Freshwater Megafauna Futures”. Gregor Kalinkat provided helpful comments on an earlier version of this article. We thank Jing Du and Karan Kakouei for their assistance with data collection.

---

**Table S1.** Summary of cover images collected from biodiversity and conservation journals

| Journal | Period of data collection | Publication frequency | Twitter account |
| --- | --- | --- | --- |
| Animal Conservation<sup>†</sup> | 2003-2016<sup>‡</sup> | Quarterly (2003-2007)<br>Bi-monthly (2008-2016) | Yes |
| Biodiversity and Conservation | 1997-2012<sup>‡</sup> | Monthly | No |
| Conservation Biology<sup>†</sup> | 1997-2016 | Bi-monthly | Yes |
| Conservation Letters<sup>†</sup> | 1998-2016<sup>‡</sup> | Bi-monthly | Yes |
| Diversity and Distributions<sup>†</sup> | 2016<sup>‡</sup> | Monthly | Yes |
| Ecography<sup>†</sup> | 2014-2016<sup>‡</sup> | Monthly | Yes |
| Global Change Biology<sup>†</sup> | 1998-2016 | 8 issues per year (1997-2001)<br>Monthly (2002-2016) | Yes |
| Journal for Nature Conservation | 2002-2016<sup>‡</sup> | Quarterly (2002-2010)<br>Bi-monthly (2011-2016) | No |
| Journal of Applied Ecology<sup>†</sup> | 1997-2016 | Bi-monthly | Yes |
| Journal of Fish and Wildlife Management | 2010-2016<sup>‡</sup> | Biannual | No |
| Northeastern Naturalist | 1997-2016 | Quarterly | No |
| Oryx<sup>†</sup> | 2007-2016 | Quarterly | Yes |
| Pachyderm | 1997-2016 | Biannual (1997-2013)<br>Annual (2014-2016) | No |
| Southeastern Naturalist | 2002-2016<sup>‡</sup> | Quarterly | No |
| Systematics and Biodiversity<sup>†</sup> | 2003-2016<sup>‡</sup> | Quarterly (2003-2014)<br>Bi-monthly (2015-2016) | No |
| The Southwestern Naturalist | 1997-2016 | Quarterly | No |
| Tropical Conservation Science | 2008-2016<sup>‡</sup> | Quarterly (Bi-monthly in 2013) | No |
| Wildlife Society Bulletin | 1997-2006; 2011-2016<sup>‡</sup> | Quarterly | No |

<sup>†</sup>These journals show Altmetric scores for each article on their websites.
<sup>‡</sup>These journals started changing cover images regularly after 1997. All cover images have been included. Wildlife Society Bulletin has been paused between 2007-2010.

## Notes

### Competing Interest Statement

The authors have declared no competing interest.

